# A Reductive Uric Acid Degradation Pathway in Anaerobic Bacteria

**DOI:** 10.1101/2025.05.19.654654

**Authors:** Zhi Li, Wei Meng, Zihan Gao, Wanli Peng, Jianhao Zhang, Yining Wang, Xiaoxia Wu, Zipeng Zhao, Chuyuan Zhang, Zhuohao Tang, Zhujun Nie, Shaohua Wu, Benjuan Wu, Yang Tong, Yiling Hu, Zehan Hu, Yifeng Wei, Yan Zhang

## Abstract

Uric acid is a key intermediate in purine degradation across diverse organisms, while its accumulation in humans leads to inflammation and gout disease. Aerobic organisms degrade uric acid via a well-known “oxidative pathway” involving dearomatization of the purine core catalyzed by uric acid oxidases or dehydrogenases. The ability to degrade uric acid is also widespread in anaerobic bacteria, including gut bacteria, although the mechanisms are incompletely understood. Here, we report the biochemical characterization of a recently identified uric acid degradation gene cluster from *Escherichia coli*, and show that it encodes a “reductive pathway” for uric acid degradation. In this pathway, uric acid is first reduced to isoxanthine (2,8-dioxopurine) by a xanthine dehydrogenase homolog (XdhD), followed by dearomatization of the purine core catalyzed by a flavin-dependent reductase (YgfK). Stepwise cleavage of the pyrimidine and imidazole rings forms 2,3-diureidopropionate, and stepwise cleavage of the 2- and 3-ureido groups then forms 2,3-diaminopropionate, which is cleaved by a pyridoxal 5’-phosphate-dependent lyase (YgeX) to pyruvate and ammonia. Detection of isoxanthine in clinical serum samples suggests it is a physiologically relevant circulating metabolite. A probiotic *E coli* Nissle strain was engineered for constitutive overexpression of the gene cluster, and oral administration in a uricase-knockout hyperuricemic mouse model significantly reduced the serum uric acid level and alleviated associated kidney injury, suggesting a potential route towards uricolytic probiotics.

## Introduction

Uric acid is a ubiquitous intermediate in purine degradation, playing important roles in the physiology of various animals. In most mammals, the enzyme uricase converts uric acid into the more water-soluble allantoin, which is readily excreted (**Figure 1a**) ^1,2^. In contrast, humans and other higher primates lack a functional uricase and instead directly eliminate uric acid, primarily through urine with a smaller contribution via the gut (**Figure 1a**) ^3,4^. In moderate levels, the electron-rich uric acid functions as a systemic antioxidant and supports neuroprotection ^1,5,6^. However, elevated uric acid levels (hyperuricemia) can lead to a range of morbidities, and are treated pharmacologically with drugs like allopurinol or benzbromarone (**Figure 1a**). Due to its poor solubility and high crystallinity, uric acid readily forms sodium urate crystals, which can deposit in the kidneys to cause kidney stones, or in joints to trigger gout, a form of inflammatory arthritis. Conversely, this low solubility is advantageous in reptiles, birds and insects, where uric acid serves as the primary nitrogenous waste and is excreted in a semi-solid form, aiding in water conservation ^7,8^.

**Figure 1.**
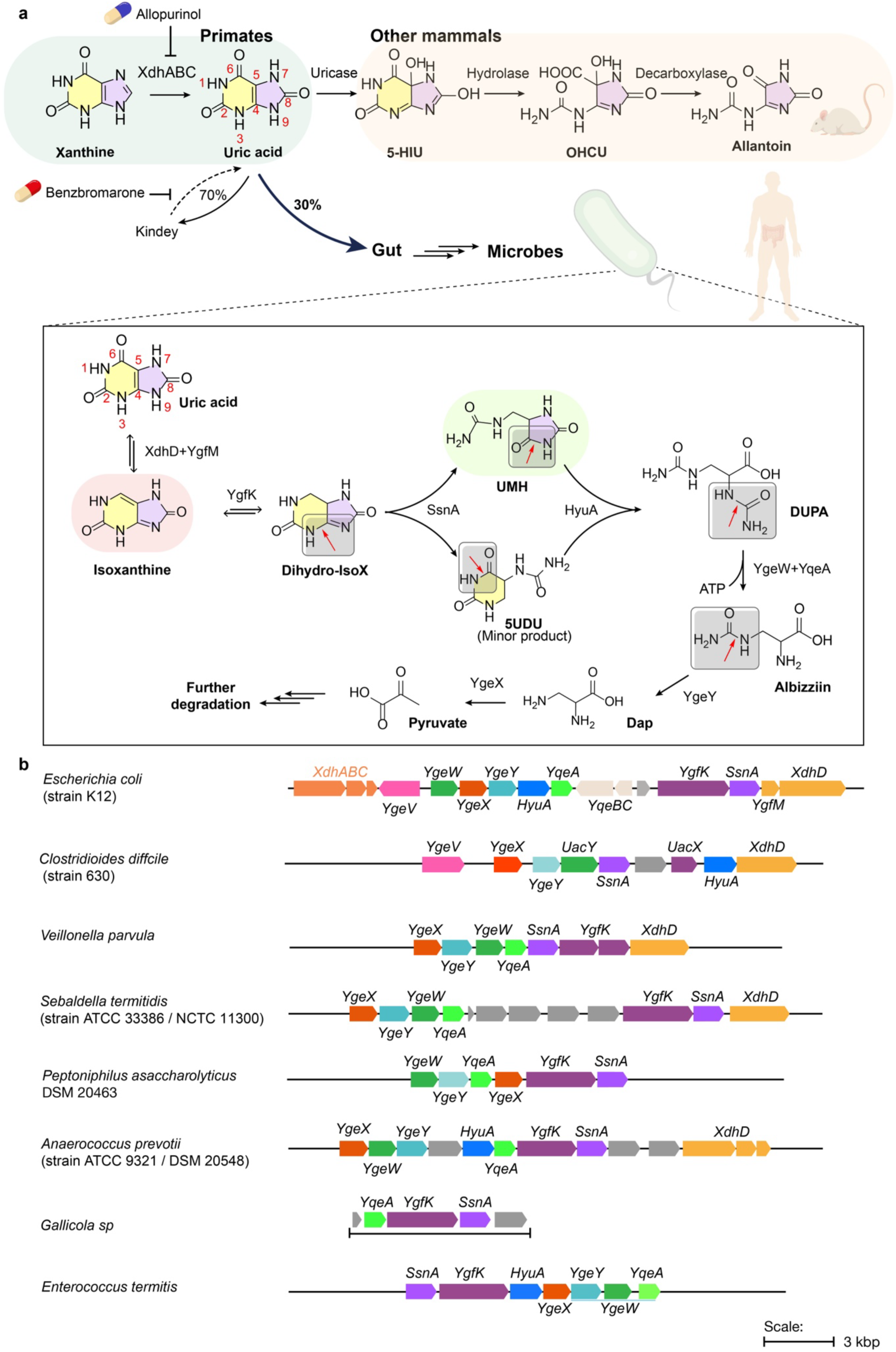
Purine degradation pathways and gene clusters. a, Purine degradation pathways, upper panel (previously known), oxidative purine degradation pathway, lower panel (boxed, this work), reductive purine degradation pathway. The physiological fate of uric acid in mammals and drugs for gout disease and drug targets are labeled. **b,** Representative bacterial gene clusters related to the reductive degradation pathway.

The study of bacterial purine and pyrimidine degradation has a long and rich history, revealing diverse strategies for cleaving the aromatic N-heterocyclic cores. For pyrimidines, these include the oxidative and reductive pathways, where ring cleavage is preceded by uracil oxidative or reductive dearomatization ^9,10^, as well as the alternative Rut ^11,12^ and URC pathways (**Extended Data** Figure 1a-d) ^13,14^. The well-known purine degradation pathway generally involves deamination of adenine and guanine to hypoxanthine and xanthine, followed by oxidation to uric acid by the molybdopterin- (MoCo-) dependent enzyme xanthine dehydrogenase (XDH) ^15,16^. In aerobic organisms, uric acid is oxidatively dearomatizated to 5- hydroxyisourate (5-HIU) by various urate oxidases and dehydrogenases^17,18^ ^19^, followed by hydrolysis and decarboxylation to form allantoin for further degradation (**Figure 1a**) ^20^.

Purine and uric acid degradation pathways that operate independently of oxygen or external electron acceptors are of particular interest due to their relevance to the anaerobic human gut microbiome. The first such purine / uric acid degradation pathway was described in *Clostridia*^21^ and elucidated by Rabinowitz *et al.*^22^. Here, XDH catalyzes the reversible reduction of uric acid to xanthine, which is then cleaved by xanthine amidohydrolase (xanthinase), followed by a series of hydrolytic and decarboxylative steps yielding glycine ^23^. The gene cluster encoding this oxygen-independent “xanthinase pathway” was only recently identified, and found to occur in *Clostridia* and *Bacilli* ^24^. While mechanistically simple, its overall rate is constrained by the slow kinetics of xanthinase (*k_cat_* =0.1 s^−1^) ^24^. Uric acid was also degraded anaerobically by diverse bacteria from the uric acid-rich termite gut, despite lacking the xanthinase pathway ^25–27^. *Streptococcus* UAD1 (now *Enterococcus termitis* UAD1) degraded uric acid when supplied with formate as a co-substrate, yielding CO_2_, NH_3_ and acetate ^28^. Urea was not detected as an intermediate, and omission of formate led to accumulation of a fluorescent metabolite in the culture medium.

The model bacterium *E. coli* oxidizes purines to uric acid via a XDH homolog (XdhA), and is also capable of degrading allantoin ^29^. However, it cannot oxidize uric acid to allantoin, thus lacking a complete pathway for aerobic purine degradation. Under anaerobic conditions, *E. coli* exhibits limited uric acid degradation in the presence of formate, previously proposed to involve two uncharacterized oxidoreductases (AegA and UacF) ^30^, located next to the uric acid-specific permease (UacT) ^31^. Recent work by the laboratories of Dodd et al. and Rey et al. provided a key advance by identifying a widely distributed gene cluster involved in anaerobic uric acid degradation, present in gut bacteria and contributing to host purine homeostasis^2,32^. This gene cluster is adjacent to XdhA and UacT in the genome of *E. coli*, and is also present in other previously characterized anaerobic uricolytic bacteria ^32,33^ (**Figure 1b**), providing a basis for further investigations into the biochemical details of this pathway. In contrast to the uricase-dependent “oxidative pathway”, we hypothesize that this gene cluster encodes a “reductive pathway” for uric acid degradation (**Figure 1a**, boxed), analogous to the reductive pathway for pyrimidine degradation (**Extended data Figure 1a**).

## Results

### Comparative analysis of *E. coli* and *C. difficile* uricolytic gene clusters

The *E. coli* uricolytic gene cluster encodes a XDH homolog (XdhD-YgfM), a flavoenzyme (YgfK), an amidohydrolase related to PydB (SsnA), D-stereospecific “phenylhydantoinase” (HyuA), a carbamoyltransferase (YgeW), a carbamate kinase (YqeA), a peptidase (YgeY), and a diaminopropionate ammonia lyase (YgeX), which is equally active on the L- and D-isomers^34^. These enzymes suggest a pathway, in which the central C3 moiety of uric acid is released as diaminopropionate, requiring a net four-electron reduction and multiple hydrolytic steps (**Figure 1a**, boxed).

Further insight was gained by comparing the *E. coli* gene cluster to that of *C. difficile*, which is also capable of degrading uric acid^33^. In *C. difficile*, YgfK is replaced by a homolog of PydA (which we designate UacX), an enzyme catalyzing reductive dearomatization of uracil in the Pyd pathway (**Extended Data** Figure 1a, **Figure 1b**). Additionally, in *C. difficile*, YgeW and YqeA are replaced by a homolog of N-acyl-D-amino acid hydrolase (D-aminoacylase), which we designate UacY (**Figure 1b**). Structural modeling reveals that its active site closely resembles that of the characterized D-aminoacylase from Bordetella (PDB: 3GIQ) with conserved residues binding the D-aminoacyl moiety (**Extended Data** Figure 2a), suggesting that YgeW and UacY catalyze the cleavage of a similar N-acyl amino acid substrate.

Structural modeling suggests that *E. coli* XdhD-YgfM form a complex resembling *Rhodobacter capsulatus* XDH (PDB: 1JRO), with XdhD containing the MoCo-binding hydroxylase/dehydroxylase site, and YgfM containing the FAD-binding site for electron transfer from NAD(P)H (**Extended Data** Figure 2b). Structural modeling also suggests *E. coli* YgfK contains a FMN-binding site likely for substrate reduction, and a FAD site likely for electron transfer from NAD(P)H, resembling the architecture of *Sus scrofa* PydA (**Extended Data** Figure 2c). By contrast, *C. difficile* XdhD lacks the YgfM subunit, and *C. difficile* UacX contains a FMN-binding site but lacks the FAD-binding domain. Instead, the architecture of *C. difficile* UacX resembles Clostridial PydA (PydAc)^35^, suggesting an alternative electron donor for both Clostridial enzymes, XdhD and UacX, such as reduced Fdx.

Variants of the gene cluster are also present in other organisms previously reported for anaerobic uric acid degradation (**Figure 1b**), including *Enterococcus faecalis* isolated from chicken cecum or human feces, *Peptoniphilus asaccharolyticus* DSM 20463, the RNA-degrading *Peptococcus prevotii* (now *Anaerococcus prevotii*) from human skin, and relatives of the obligate purine-fermenting *Peptostreptococcus barnesae* (now *Gallicola barnesae*) from chicken feces.

Together, the composition of the two gene clusters offers key insights into the key steps involved in the proposed pathway (**Figure 1a**, boxed). The net four-electron reduction likely involves a dehydroxylation step catalyzed by XdhD, and a reductive dearomatization step catalyzed by YgfK / UacX. Following dearomatization, the two rings are likely cleaved by SsnA and HyuA. The two ureido groups linked to the diaminopropionate moiety are likely cleaved by YgeY and YgeW / UacY, with YgeY cleaving the 3-ureido group, and YgeW / UacY cleaving the 2-ureido group, as suggested by the substrate specificity of D-aminoacylases.

### XdhD catalyzes reduction of uric acid to 2,8-dioxopurine (isoX)

*Breznak et al.* reported that in the absence of formate as a co-substrate, Enterococcus UAD1 degrades uric acid incompletely, secreting a fluorescent compound with excitation and emission maxima at 308 and 379 nm (pH 7.0)^36^. To identify potential intermediates in the pathway, we screened a panel of aromatic metabolites, including uric acid, hypoxanthine, xanthine, 2,8-dioxopurine (isoxanthine, isoX), 2,6-dioxopurine, 5-aminouracil, and 5- ureidouracil, and found that only isoX displayed a closely matching fluorescence emission peak at 363 nm upon excitation at 308 nm (**Extended Figure 3a-b**). We therefore hypothesized that isoX is an intermediate, formed via dehydroxylation of uric acid catalyzed by XdhD.

To test this hypothesis, *E. coli* XdhD was expressed with a C-terminal ProtA affinity tag from the low-copy plasmid *p*SC101, to prevent overwhelming the cofactor biosynthetic pathway. Affinity purification of XdhD showed co-purification with YgfM (**Figure 2a**). Incubation of the purified protein with uric acid and NADH resulted in the formation of isoX as the major product, as identified by LC-MS and co-elution with a commercial standard (**Figure 2b**). Boltz- 1 prediction of XdhD in complex with uric acid and structural comparison reveal that E606 present in the active site of XdhD but absent in XdhA may contribute to the orientation of the substrate and formation of isoX (**Figure 2c**).

**Figure 2.**
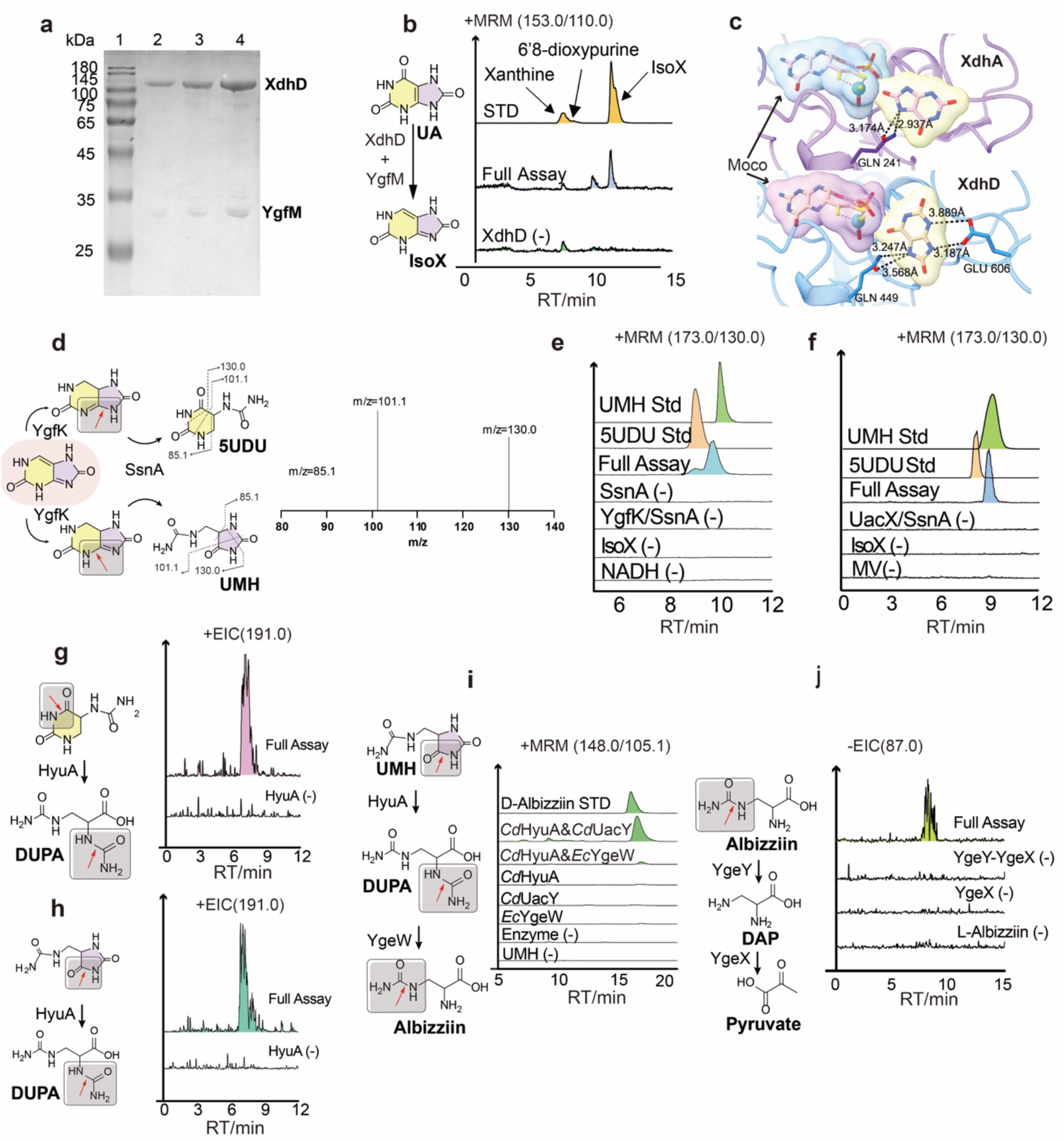
Stepwise *In-vitro* reconstitution of the reductive uric acid degradation pathway. a, SDS-PAGE of moderately overexpressed *Ec*XdhD-YgfM. **b,** Assays on *Ec*XdhD-YgfM. LC-MRM-MS. **c,** Comparison of the activity site and substrate orientations in *Ec*XdhA and *Ec*XdhD. The key Glu606 residue in *Ec*XdhD but absence in *Ec*XdhA is labeled. **d-f,** Assays on recombinantly produced reconstituted YgfK-SsnA. LC-MRM-MS (173.0/130.0 transition) chromatogram monitoring product formation in the YgfK and SsnA coupling assay. **g-h,** Assays on recombinant *Cd*HyuA. LC-MS extracted ion (m/z 191.0) chromatographs monitoring product formation in the *Cd*HyuA assay. **i,** Assays on recombinant *Cd*HyuA- *Cd*UacY. LC-MRM-MS (148.0/105.1 transition) chromatogram monitoring product formation in the *Cd*HyuA and *Cd*UacY coupling assay. **j,** Assays on recombinant YgeY-YgeX. LC-MS extracted ion (m/z 87.0) chromatographs monitoring product formation in the YgeY-YgeX coupling assays.

**Figure 3.**
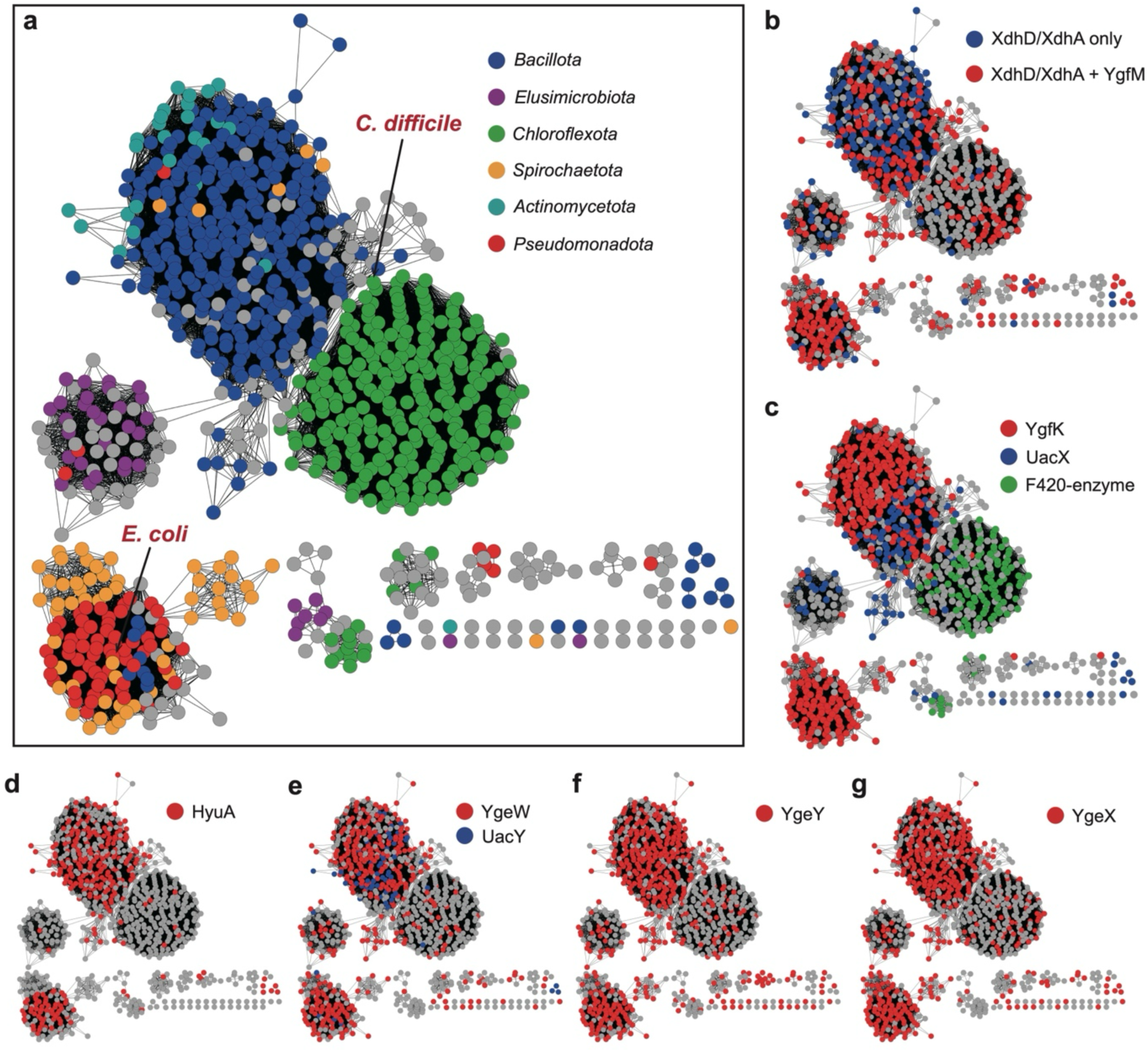
Sequence Similarity Network of the SsnA family (IPR017700) in bacteria. a, Nodes colored according to bacterial phyla: Blue (*Bacillota*), Red (*Pseudomonadota*), Green (*Chloroflexota*), Purple (*Elusimicrobiota*), Orange (*Spirochaetota*), Teal (*Actinomycetota*). **b-g**, Nodes colored according to the presence of specific protein families of the different uricolytic enzymes within a 15-gene neighborhood. **b** XdhD or XdhA (PF01315) with or without YgfM (PF00941). **c** YgfK (IPR017701), UacX (PF01180), or putative F420-dependent reductase (PF00296). **d** HyuA (IPR011778). **e** YgeW (PF00185) or UacY (PF07969). **f** YgeY (PF01546). **g** YgeX (PF00291).

### YgfK and SsnA catalyze reductive dearomatization and ring cleavage of isoX

The remaining enzymes in the pathway were individually overexpressed and purified (**Extended Data** Figure 4). The YgfK 4Fe-4S clusters were reconstituted anaerobically and characterized by UV-Vis and EPR spectroscopy^37,38^ (**Extended Data** Figure 5a-b). Activity assays of individual enzymes with isoX showed no detectable activity. However, anaerobic incubation of isoX with YgfK, SsnA, and NAD(P)H resulted in the formation of ureidomethyl- hydantoin (UMH) as the major product and 5-ureido-dihydrouracil (5UDU) as a minor product, as identified by LC-MS and co-elution with commercial standards (**Figure 2d-f**). These results are consistent with reductive dearomatization of isoX by YgfK, followed by hydrolytic cleavage by SsnA. Failure to detect the reduction product with YgfK alone may reflect the unfavorable thermodynamics of this reaction, which requires coupling to hydrolysis by SsnA, consistent with the co-localization of the two genes in many organisms (**Figure 1b**). A possible catalytic mechanism for formation of the two SsnA products is shown in **Extended Data** Figure 5c.

### Enzymes catalyzing cleavage of ureidomethyl-hydantoin (UMH) to diaminopropionate

Given the structural similarity of UMH and 5UDU to known HyuA substrates, we assayed HyuA activity using the commercially available racemic compounds, and confirmed cleavage of both by LC-MS (**Figure 2g-h**). Incubation of UMH with HyuA, in the presence of either UacY or YgeW and phosphate, resulted in the formation of 3-ureido-2-aminopropionate (albizziin), as confirmed by LC-MS and co-elution with a commercial standard (**Figure 2i**). The reaction peak with YgeW was notably smaller than that with *Cd*UacY, likely reflecting the less favorable thermodynamics of the phosphorylysis reaction. Finally, incubation of albizziin with YgeY and YgeX yielded pyruvate as the terminal product of the pathway (**Figure 2j**).

### Variants of the uricolytic gene cluster in different bacteria

SsnA is the most identifiable of the uricolytic enzymes, catalyzing a key step in the pathway and serving as a robust marker for identifying uricolytic gene clusters. To explore the distribution of uricolytic gene clusters across bacteria, we constructed a Sequence Similarity Network (SSN) of the SsnA family (IPR017700). SsnA is prevalent in Pseudomonadota (e.g., *E. coli*), Bacillota (e.g., *C. difficile*), as well as Chloroflexota and Actinomycetota. It is also present in Spirochaetota and Elusimicrobiota, two phyla commonly found in the termite gut (**Figure 3a**). Genome neighborhood analysis within a 15-gene window revealed that SsnA colocalizes with the upstream pathway genes. These include XdhA/XdhD homologs, some containing the YgfM FAD-binding domain (**Figure 3b**), and either YgfK (as in *E. coli*) or UacX (as in *C. difficile*) (**Figure 3c**). Notably, Chloroflexota lack these enzymes and instead contain a putative F420-dependent dehydrogenase belonging to the “luciferase family”, suggesting an alternative mechanism for isoX reduction (**Figure 3c**). In many organisms, SsnA also colocalizes with the downstream pathway genes HyuA (**Figure 3d**), YgeW/UacY(**Figure 3e**), YgeY (**Figure 3f**), and YgeX(**Figure 3g**). These findings suggest the widespreadness of the uricolytic pathway in different bacterial lineages.

### An engineered *E. coli* strain CBT2.0 overexpressing the uricolytic gene cluster

Recent studies by Dodd *et al*. demonstrated that microbiota depletion in uricase-deficient mice leads to hyperuricemia, and that treatment with antibiotics targeting anaerobic microbiota increases the risk of gout in humans^2^. Rey *et al*. further showed that colonizing gnotobiotic mice with purine-degrading bacteria modulates systemic levels of uric acid, and that uricolytic bacteria harboring this gene cluster are correlated with serum uric acid levels in humans^32^. By extension, we hypothesized that engineering gut bacteria for constitutive overexpression of the uricolytic pathway could enhance uricolytic activity by the gut microbiome, and ensuring consistent pathway activation and bypassing repression by energy-rich dietary molecules such as glucose.

Starting with the parental *E. coli strain EcNc*, derived from the probiotic *E. coli Nissle 1917* (*EcN*) by ejection of two cryptic plasmids pMUT1 and pMUT2, CRISPR-Cas9 genome editing was used to insert a hybrid promoter cassette comprising (1) a strong constitutive *gapA* promoter and (2) an anaerobic nirB promoter into the intergenic region between ygeV and ygeW, replacing the native Crp (cAMP Receptor Protein)-binding site (**Figure 4a**). The resulting strain was named CarBT4gout_2.0 (**CBT2.0, Figure 4a**), and overexpression of uricolytic genes was confirmed by RNA-seq analysis (**Extended Data** Figure 6). For more detailed biochemical analysis, the Δ*xdhA* and Δ*xdhD* deletion mutants were generated for *E. coli* strains MG1655 strains (**Figure 4 b-c**) and CBT 2.0 strains (**Figure 4 d-e**). Δ*ygfK* deletion mutants were also generated for *EcN* (**Figure 4 f-k**).

**Figure 4.**
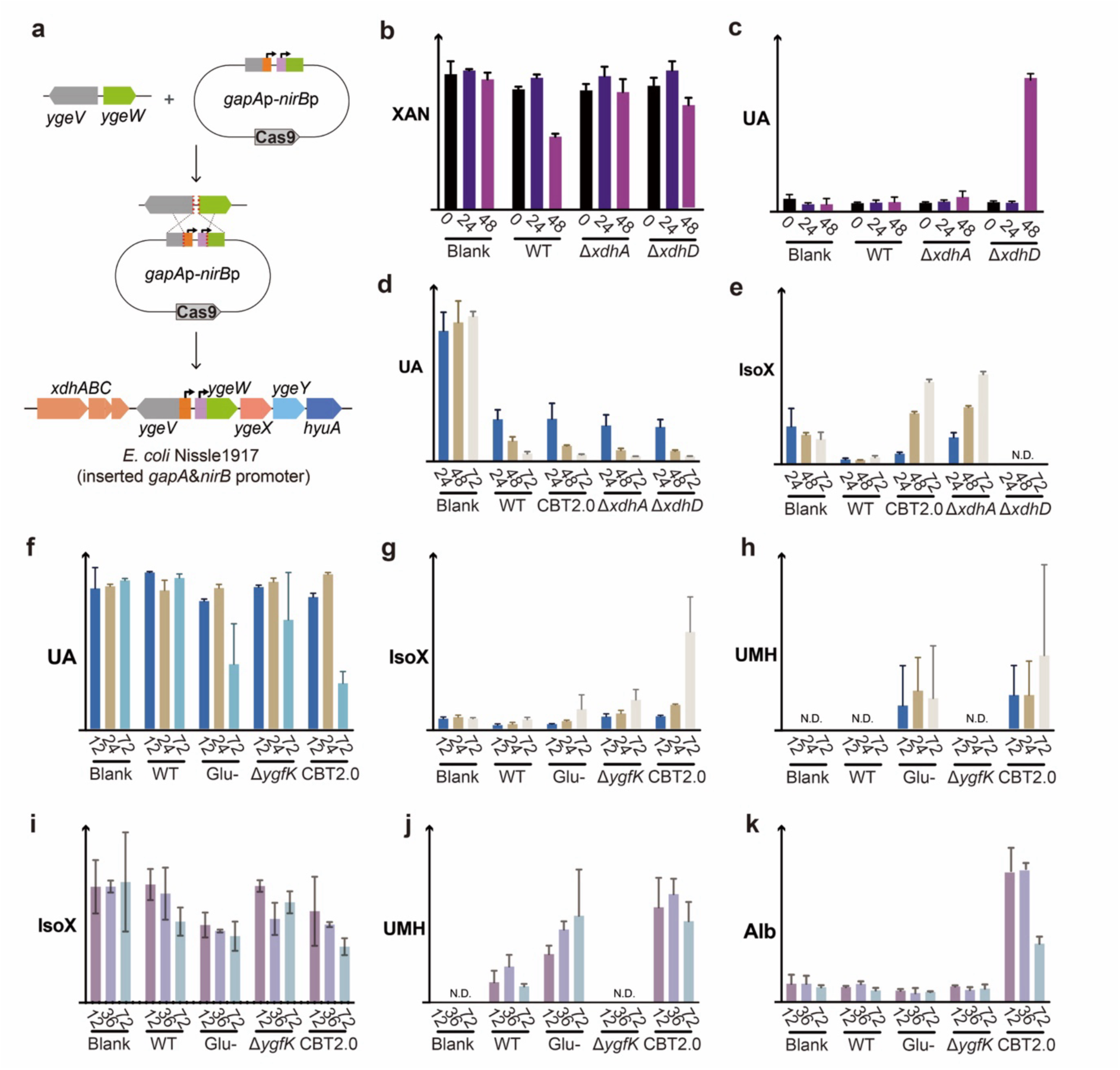
Anaerobic purine metabolic intermediates in *E. coli*. a, Design of CarBT4gout 2.0. **b-c,** Xanthine consumption and UA accumulation in various *E. coli* strains grown anaerobically in minimal media. **d-e,** Uric acid consumption and isoxanthine accumulation in various *E. coli* strains grown anaerobically in LB media. **f-h,** Uric acid consumption and metabolic intermediate accumulation in various *E. coli* strains grown anaerobically in LB media. **i-k,** IsoX consumption and metabolic intermediate accumulation in various *E. coli* strains grown anaerobically in LB media.

Dodd *et al*. previously showed that gut bacteria metabolize uric acid via two distinct pathways: reduction to xanthine (via XdhA) or uricolysis (via SsnA, YgfK etc.)^2^. To evaluate the proposed uricolytic pathway *in vivo*, we cultured selected mutant strains with the various purine compounds, collected culture supernatants at defined time points, and analyzed them by LC-MS. For cultures in M9 glucose medium supplemented with xanthine, both WT and *ΔxdhD* consumed xanthine, while *ΔxdhA* did not, consistent with the role of XdhA in xanthine oxidation (**Figure 4b**). In *ΔxdhD*, xanthine consumption was accompanied by uric acid accumulation, consistent with the role of XdhD in subsequent uric acid degradation (**Figure 4c**).

For cultures with LB supplemented with uric acid, all strains tested showed uric acid degradation, likely via XdhA or XdhD (**Figure 4d**). IsoX was detected at low levels in fresh LB medium, and further accumulated only in CBT and *ΔxdhA*, suggesting increased flux through XdhD in these strains (**Figure 4e**). Consistently, no isoX was observed in *ΔxdhD* (**Figure 4e**). For cultures with LB supplemented with uric acid and glucose, uric acid degradation was repressed in WT but not in CBT, indicating that CBT can bypass glucose repression (**Figure 4f**). IsoX accumulated in CBT (**Figure 4g**), similar to observations in LB without glucose (**Figure 4e**). UMH was detected only in CBT and in WT grown without glucose, conditions where uric acid degradation occurred, supporting its role as a downstream intermediate (**Figure 4h**). For cultures with LB supplemented with isoX and glucose, all strains tested showed isoX degradation, but degradation was faster in CBT and in WT grown without glucose (**Figure 4i**), mirroring trends with uric acid (**Figure 4f**). UMH was detected for all strains except *ΔygfK*, consistent with the role of YgfK in converting isoX to UMH (**Figure 4h, j**). Albizziin was detected at low levels in fresh LB medium, and further accumulated only in CBT, possibly reflecting enhanced activity of upstream enzymes such as YgeW (**Figure 4k**). Overall, the detection of isoX, UMH, and albizziin supports their roles as intermediates in uric acid degradation, while the phenotypes of *ΔxdhA*, *ΔxdhD*, and *ΔygfK* validate their proposed functions. Additionally, the CBT strain demonstrates resistance to glucose repression and increased flux through the uricolytic pathway.

### *E. coli* CBT2.0 lowers uric acid levels in a hyperuricemic mouse model

To assess the uric acid-lowering potential of CBT2.0 in a host organism, we utilized a uricase-deficient (UOX⁻/⁻) mouse model of hyperuricemia. Mice were randomly assigned to three groups, and received daily intragastric gavage of CBT2.0, wild-type *EcN* (WT), or PBS vehicle control for 6 weeks, with weekly blood collection to monitor plasma levels of uric acid (UA), urea nitrogen (UN) and creatinine (CRE) (**Figure 5a**). Throughout the treatment period, mice administered CBT2.0 exhibited a significant reduction in plasma UA levels compared to the PBS control group. By week 6, the CBT2.0 group showed a mean UA concentration of 171.63 ± 91.59μmol/L, whereas the PBS group maintained elevated levels at 463.26 ± 70.81μmol/L (*p* < 0.001) (**Figure 5b**). In contrast, the WT group showed a more modest reduction in UA levels compared to PBS at certain time points, but not consistently throughout the entire study period (**Figure 5b**). These findings indicate that, compared to WT, CBT2.0 more reliably lowers systemic uric acid levels in UOX⁻/⁻ mice.

**Figure 5.**
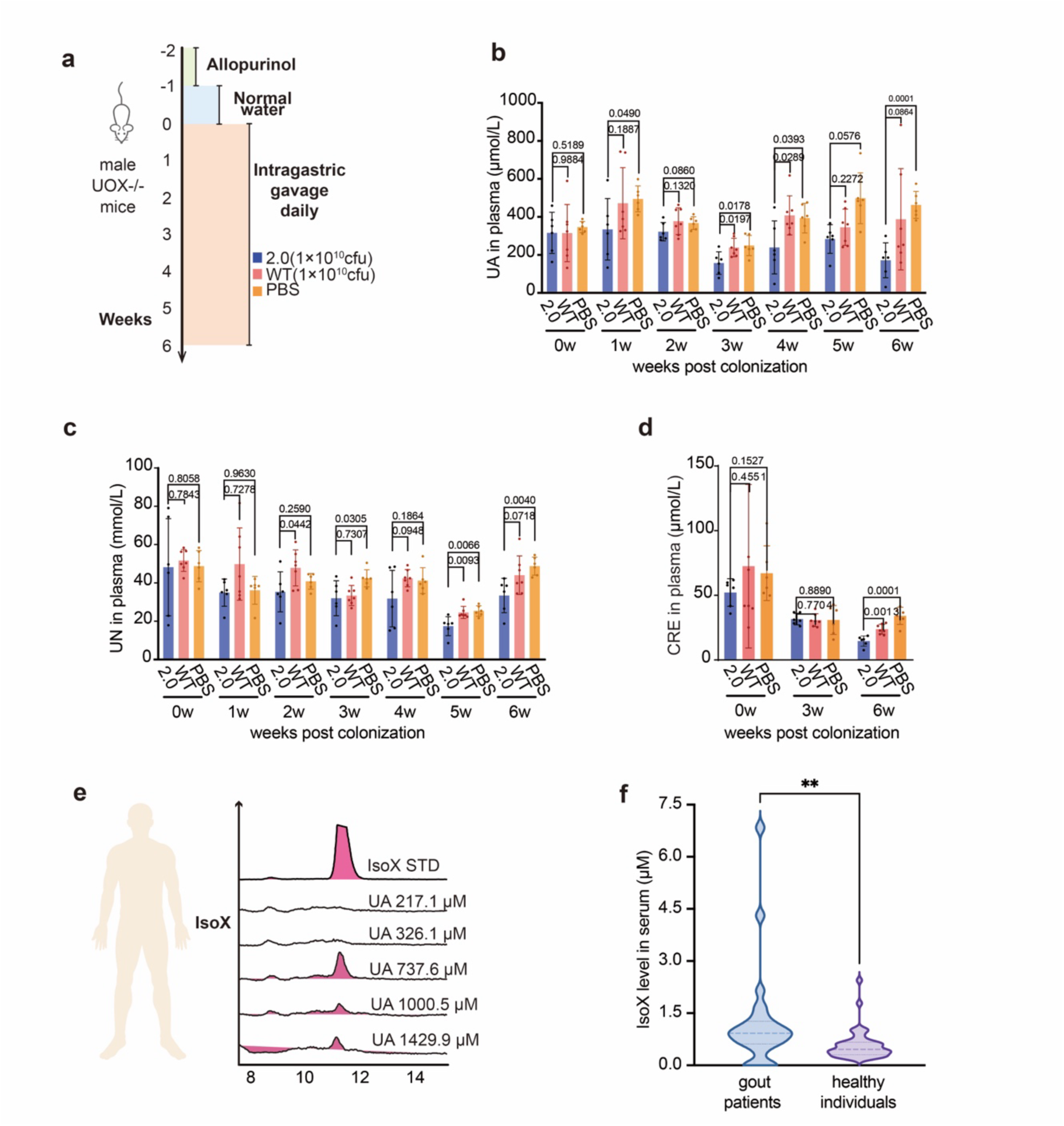
Physiological and pathological relevance of isoxanthine in human plasma and the effectiveness of engineered probiotics in an animal gout model. **a**, Mouse experimental design. Male uricase-deficient (UOX-/-) mice were withdrawn from allopurinol for one week at week -1 and then orally gavaged once daily for six consecutive weeks starting from week 0 with CarBT4gout 2.0 (2.0, blue, 1 × 10^10^ CFU), wild-type *Escherichia coli* Nissle 1917 (WT, red, 1 × 10^10^ CFU), or vehicle control (PBS, orange, 200 μL). Plasma was collected weekly. **b**, Plasma uric acid (UA) levels in mice gavaged with 2.0, WT, or PBS. **c**, Plasma urea nitrogen (BUN) levels in mice gavaged with 2.0, WT, or PBS. **d,** Plasma creatinine (CRE) levels in mice gavaged with 2.0, WT, or PBS at week 6. All shown data in bar represent mean ± SD from n = 6-7 mice per group; individual values are overlaid. Statistical comparisons were performed with two-tailed unpaired *t*-tests; exact *p*-values are indicated above brackets. **e-f**, IsoX was detected across human plasma samples with different uric acid levels.

The physiological effects of hyperuricemia in mice differ somewhat from those in humans, with urate crystals more prone to depositing in the kidneys rather than joints, leading to potentially fatal acute kidney injury. Impaired renal excretory function typically results in systemic accumulation of CRE and UN. To evaluate the renal effects of three treatments, we monitored plasma UN throughout the study and assessed CRE levels at weeks 0, 3, and 6. By week 6, the CBT2.0 group exhibited significantly lower UN levels (33.40 ± 8.96 mmol/L) compared to the WT group (44.02 ± 10.07 mmol/L, *p* = 0.0718) and PBS group (48.75 ± 4.70 mmol/L, *p* = 0.004) (**Figure 5c**). A parallel trend was observed in CRE levels, with the 2.0 group demonstrating markedly reduced level (14.61 ± 3.97 μmol/L) relative to the PBS group (34.14 ± 6.66 μmol/L, *p*< 0.0001) and WT group (23.86 ± 3.81 μmol/L, *p* = 0.0013) (**Figure 5d**). Notably, CBT2.0-treated mice displayed lower UA levels than the PBS group as early as week 1 (although statistical significance was not reached at weeks 2 and 5, *p* > 0.05) (**Figure 5b**), and a significant BUN reduction in the 2.0 group compared to PBS controls was first observed at week 3 (**Figure 5c**), while no significant difference in CRE levels was detected at this time point (**Figure 5d**), which suggest that the renal protective effects of the CBT2.0 treatment, potentially mediated through UA reduction, may exhibit a time-lagged progression.

### Detection of isoX in clinical serum samples

The detection of isoX secretion in *E. coli* and previously in Enterococcus sp.^36^ prompted us to investigate whether it is also detectable in serum and whether it correlates with uric acid levels. Using our in-house LC-MRM-MS assay, the newly discovered metabolite isoX was sensitively quantified in serum via the optimized transition m/z 153.0 /110.0. A preliminary analysis of human serum samples with known uric acid levels suggested that isoX levels could be used to distinguish three diagnosed gout patients from two healthy controls (**Figure 5e**). In a subsequent analysis of 68 clinical human serum specimens (**Figure 5f**), IsoX serum levels in 25 patients with gout were significantly elevated compared to 43 non-gout controls, with a statistically significant difference between the two groups (*p* < 0.01). This suggests that isoX may serve as a biomarker, warranting further investigation.

## Discussion

Characterization of the enzymes encoded by a widely distributed anaerobic uric acid degradation gene cluster in *E. coli* reveals a “reductive pathway” for purine catabolism. All three purine degradation pathways studied to date initiate ring cleavage at the pyrimidine moiety, but employ different strategies to overcome the aromatic stabilization. In the “xanthinase pathway,” annelation to the aromatic imidazole ring diminishes the aromatic stabilization of the uracil moiety of xanthine, thereby permitting direct hydrolytic cleavage, albeit with slow kinetics^24^. Conversely, in the “reductive pathway,” the imidazole ring is dearomatized due to the 8-oxo group in isoxanthine, which facilitates subsequent reductive dearomatization of the uracil moiety and subsequent cleavage, in a manner directly analogous to PydA and PydB in the reductive pyrimidine degradation (Pyd) pathway.

The molecular details of this pathway have important implications for understanding purine and uric acid metabolism in the anaerobic human gut microbiota, with potential clinical implications relevant to hyperuricemia and gout. While early studies focused on uricolytic bacteria from insects and birds, which excrete large amounts of uric acid, gut excretion is also a significant route of uric acid elimination in humans and other primates. Recent work by the laboratories of Dodd and Rey demonstrated that this gene cluster is broadly distributed among phylogenetically diverse human gut bacteria and can influence host purine and uric acid homeostasis^2,32^. A notable observation was the detection of isoxanthine in human plasma samples. Isoxanthine was released by *E. coli* grown on uric acid, and likely corresponds to an unidentified fluorescent metabolite previously observed in *Enterococcus* sp. grown on uric acid in the absence of formate (Breznak et al. 1980)^36^, showing that anaerobic bacterial purine metabolism can produce circulating metabolites, as previously reported for aromatic amino acid metabolism^39^. Given its strong fluorescence, isoxanthine merits further investigation as a potential biomarker for purine metabolic disorders, presumably more advantageous and convenient than enzyme-needed uric acid measurement.

A widely proposed goal is to harness microbiome-based strategies to modulate uric acid cycling, yet the regulation of uric acid metabolism genes and the metabolic flux of uric acid degradation across diverse gut bacteria remain poorly understood, particularly within the chemically complex environment of the digestive tract. Our experiments with an engineered strain of the probiotic *E. coli* Nissle in a uricase-deficient mouse gout model showed that constitutive expression of the uric acid degradation gene cluster led to reduced host plasma uric acid levels, supporting the potential of engineered probiotics for managing hyperuricemia. For this application, the “xanthinase pathway” is attractive for its simplicity but is constrained by the slow kinetics of xanthinase, while the “reductive pathway” is more widespread but relies on complex cofactors like Mo-Co, Se, and 4Fe-4S clusters.

Bioinformatic analyses reveal the existence of multiple variants of the “reductive pathway” enzymes, which parallel earlier observations in the Pyd pathway, with the *E. coli* XdhD and YgfK relying on NADH as the reductant, the corresponding *C. difficile* XdhD and UacX likely relying on Fdx as the reductant, and gene cluster in Chloroflexota bacteria containing a putative isoX reductase relying on coenzyme F420. Another key difference is the cleavage of the 2 ureido group through an irreversible hydrolytic reaction by *C. difficile* UacY, and reversible phosphorylytic reaction by *E. coli* YgeW, forming carbamoyl phosphate to support ATP generation. These pathway variants may reflect differences in redox mediators and energy conservation strategies. Further studies are needed to determine the optimal combination of alternative enzymes for engineering uricolytic probiotics, balancing uric acid flux and energy conservation.

## Methods

### Strains, cultures, and chemicals

The strains and plasmids used are listed in **Table S1**. For uric acid utilization assays, *Escherichia coli* cells were cultured anaerobically at 37 °C in M9 minimal medium (1×M9 salts, 2 mM MgSO₄, 0.1 mM CaCl₂, and 4 g/L glucose, pH 7.0) in an anaerobic vial, unless otherwise stated. For genetic manipulations, cells were grown aerobically in LB broth at 37 °C. For oxygen-free cultivation, the medium was prepared in a Lab2000 glovebox (Etelux) under an atmosphere of N₂ containing less than 5 ppm O_2_ by dissolving medium powder in degassed ddH_2_O, followed by filtration through a 0.2 μm filter. For bacteria used in mouse experiments, cells were inoculated into 5 mL of LB broth and cultured overnight at 37 °C with shaking. Cell growth in cultures were monitored by measuring optical density at 600 nm (OD_600_). All chemicals and reagents were of analytical grade and are detailed in **Table S2**.

### Standard DNA manipulation

All oligonucleotides in this study were synthesized by Sangon Ltd and listed in **Table S3**. DNA polymerases, T4 DNA ligases, and all restriction endonucleases were obtained from TaKaRa Bio Inc and ThermoFisher Scientific Inc. Genomic DNAs and plasmid DNAs were extracted using the TIANamp Bacteria DNA Kit and the TIANprep Mini Plasmid Kit, respectively (Tiangen), while amplified PCR fragments were purified using the TIANquick Midi Purification Kit (Tiangen).

Genes from *E. coli* MG1655 encoding YgeW, YgeX, YgeY, HyuA, YqeA, YgfK and SsnA (Uniprot ID: Q46803, P66899, P65807, Q46806, Q46807, Q46811 and Q46812) were amplified by colony PCR and individually inserted into the pET28a-6*His-TEV (HT) vector^40^. Gene syntheses for *Clostridium difficile* UacX (Uniprot ID: Q181U4), HyuA (Uniprot ID: Q181U3) and UacY (Uniprot ID: Q181U6), and expression were customized and inserted into the HT vector (Beijing Azenta Biotechnology Co., Ltd., China). The *xdhD* gene was cloned and expressed under its native promoters using the low-copy-number plasmid pSC101 (5-10 copies per cell) to minimize metabolic burden on the cofactor synthesis. The construct was generated by PCR amplification of the *xdhD* gene with its native promoter, followed by insertion of a TEV protease recognition site, a (GGGGS)*3 flexible linker, and a tandem Protein A (ProtA) tag at the C-terminus to facilitate affinity purification and activity assays^38^. Knockout mutants were generated via CRISPR-aided homologous recombination strategy and confirmed by sequencing ^37,41^.

### Protein expression and purification

Typically, expression plasmids were transformed into *E. coli* BL21 (DE3) cells followed by induction with 0.3 mM isopropyl β-D-1-thiogalactopyranoside (IPTG) at 18 °C for 16 h. Cell pellets were harvested and resuspended in lysis buffer (50 mM Tris-HCl pH 8.0, 200 mM KCl, 1 mM PMSF, and 1 μL DNase I). and lysed using a pre-chilled high-pressure homogenizer. Cell debris was removed by centrifugation at 10,000 × *g* for 10 min at 4 °C. Streptomycin sulfate (final concentration, 2% (w/v)) was added to the cell lysate followed by centrifugation to remove DNA. The supernatant was then applied to a 5 mL TALON Co²⁺ column (Cytiva) pre-equilibrated with buffer A [20 mM Tris/HCl, pH 7.5, 200 mM KCl, and 5 mM β- mercaptoethanol (BME)]. The column was washed with the same buffer and His-tagged proteins of interest were eluted with buffer B [20 mM Tris/HCl, pH 7.5, 200 mM KCl, and 5 mM β-mercaptoethanol (BME), 150 mM imidazole], dialyzed against buffer A at 4 °C for 3 hours to remove imidazole, and concentrated using Amicon Ultra-15 centrifugal filters (Millipore). The purified proteins were aliquoted and flash frozen in liquid nitrogen with 20 % glycerol and stored at -80 °C.

For XdhD (Q46814) expression and purification, IPTG induction was not necessary and skipped. Cells from an overnight culture were lysed followed by streptomycin sulfate precipitation. After centrifugation at 25,000 × *g* for 60 min at 4 °C, the clarified supernatant was applied to a 1 mL pre-equilibrated Rabbit IgG Beads 4FF column (Smart-Lifesciences Bio). The column was washed with 50 column volumes of lysis buffer, and on-column TEV protease cleavage (1 mg/mL) was performed for 20 h at 4 °C to remove the ProtA tag. Finally, the protein was further purified using a HiLoad 16/600 Superdex 200 prep grade column equilibrated with size-exclusion chromatography buffer (20 mM Tris-HCl, pH 8.0, 200 mM KCl, 1 mM dithiothreitol (DTT)). The purified protein was then concentrated to 100 μL and stored at - 80 °C for subsequent enzymatic assays.

### Reconstitution, UV-Vis and EPR spectroscopic characterization of YgfK Fe-S clusters

Reconstitution of iron-sulfur (Fe-S) clusters, and UV-visible spectroscopic analyses were performed as previously described²^,^³. Briefly, protein solutions were degassed under argon using a Schlenk line and subsequently transferred into an anaerobic glovebox. Reconstitution was carried out in a buffer containing 100 mM Tris-HCl (pH 7.5), 10 mM DTT, and 12 equivalents each of ferrous ammonium sulfate [(NH₄)₂Fe(SO₄)₂] and sodium sulfide (Na₂S·9H₂O). The mixtures were incubated at 4 °C overnight followed by addition of 4 equivalents of EDTA, and removal of excess reagents by buffer exchange into 20 mM Tris-HCl (pH 7.5), 200 mM KCl. UV-visible absorption spectra (220 to 800 nm) of reconstituted *Ec*YgfK were recorded using a Nanophotometer NP80 Mobile (Implen, Germany). Samples were prepared at 10 μM in anaerobic conditions and transferred into septum-sealed quartz cuvettes. For reduced spectra, 10 equivalents of Ti (III) citrate were added, and incubated for 10 min prior to measurement. EPR samples were prepared as previously described^38,42^. EPR spectra were acquired using a Bruker EMXPLUS-10/12 spectrometer. Measurements were performed at 6 K with the following instrument settings: center field, 3450.00 G; sweep width, 600.0 G; modulation frequency, 100 kHz; microwave frequency, 9.656750 GHz; modulation amplitude, 200.0 G. Microwave power was varied (1.002 mW, 5.024 mW, 10.02 mW, and 20.00 mW) using attenuation settings of 23.0 dB, 16.0 dB, 13.0 dB, and 10.0 dB, respectively. The g-factor was calibrated at 2.000.

### *In vitro* enzymatic activity assays

To assess the catalytic activities of *Ec*XdhD-YgfM and *Cc*XDH in the conversion of uric acid to isoxanthine, *in vitro* assays were conducted using freshly prepared uric acid (pH 7.5). Reactions were carried out in 20 mM Tris-HCl (pH 7.5) containing 200 mM KCl, 1 mM NADH, 1 mM uric acid and 10 μM enzyme at room temperature (RT).

For the YgeX/YgeY coupling assays, a reaction mixture of 5 mM L/D-albizziin, 1 mM CoCl₂, and 10 μM YgeX and YgeY in 20 mM Tris-HCl (pH 7.5), 100 mM KCl was incubated at 30 °C for 60 min. For the *Cd*HyuA or *Cd*HyuA and *Cd*UacY coupling assays, 20 mM Tris-HCl (pH 7.5), 100 mM KCl, 10 mM DOI-MU or 5UDU, 1 mM ZnCl₂, and 10 μM enzyme(s) were mixed and incubated at RT for 1 h. For the *Cd*HyuA and *Ec*YgeW coupling assays, 50 mM PBS (pH 7.2), 100 mM KCl, 10 mM DOI-MU or 5UDU, 1 mM MgCl₂, and 10 μM enzyme(s) were mixed and incubated at RT for 1 h.

Assays involving YgfK/UacX and SsnA coupling reaction were performed in an anaerobic glovebox. For YgfK-SsnA coupling assays, reconstituted YgfK (10 μM each) and degassed SsnA were mixed with 1 mM isoxanthine, 1 mM ZnCl₂, 1 mM NADH in 20 mM Tris-HCl (pH 7.5) and 200 mM KCl and incubated at RT for 1 h. For UacX and SsnA coupling assays, typical enzymatic assays were performed in 200 μL reaction volumes containing 20 mM Tris- HCl (pH 7.5), 200 mM KCl, 0.5 mM reduced methylviologen radical (MV•^+^), 0.35 mM flavin mononucleotide (FMN), 1 mM IsoX, 10 μM and 16.8 μM reconstituted UacX. Reactions were incubated at 4 °C for 2 h. Methylviologen (MV^2+^) was reduced to MV•^+^ using ∼1.2 equivalents of titanium (III) citrate in an anaerobic glovebox. The concentration of MV•^+^ was determined spectrophotometrically at 600 nm using an extinction coefficient (ε_600_) of 13,700 M^-^¹cm^-^¹. Controls omitting individual component either enzyme or substrate were included. All reactions were quenched by adding an equal volume of acetonitrile to precipitate proteins. Supernatants were collected after centrifugation and analyzed by LC-MS for product quantification.

### LS-MS analysis

Enzymatic reaction products were measured on an Agilent 1260 HPLC system coupled to an Agilent 6420 triple-quadrupole mass spectrometer. Separation was achieved on a ZIC-HILIC column (150 × 4.6 mm, 5 µm, 200 Å; Merck) using a linear gradient from 90 % to 50 % solvent B over 20 minutes at 0.75 mL min^-^¹. The mass spectrometer ran in positive-ion electrospray, multiple-reaction-monitoring mode (source temperature 330 °C; drying-gas flow 10 L min^-^¹; nebuliser pressure 45 psi). Monitored transitions and collision energies were: uric acid (169.0 / 141.0, 15 eV), isoxanthine (153.0 / 110.0, 15 eV), UMH and 5UDU (173.0 / 130.0, 10 eV), and D/L-Albizzin (148.0 / 105.1, 5 eV). Data were processed with Agilent MassHunter Qualitative Analysis software. HyuA, YgeX-YgeY, HyuA-YgeW assay products were analyzed in full-scan mode.

### Growth conditions and sample preparation for metabolic intermediate analysis

*E. coli* cells were inoculated into 5 mL of LB medium and cultured overnight at 37 °C with shaking. Cells were harvested by centrifugation at 4,000 × *g* for 10 min at 4 °C, washed with sterile PBS, and resuspended in oxygen-free LB medium. Cell suspensions were diluted to an OD₆₀₀ of 0.05 in 5 mL of oxygen-free LB, supplemented with 1 mM uric acid or isoxanthine, with or without 0.4% (w/v) glucose. Where indicated, 50 mM sodium formate (HCOONa) and 50 mM sodium bicarbonate (NaHCO₃) were added. At specified time points (0, 12, 24, 48, and 72 h), culture density (OD₆₀₀) was measured, and concentrations of uric acid, isoxanthine, DOI- MU and D/L albizziin were determined by LC-MS described above using MRM mode. Data represent the mean ± standard error from three independent biological replicates.

### RNA Extraction and Sequencing

Cultures of CarBT4gout 2.0 and EcN1917 wild-type strains were grown anaerobically in LB medium at 37 °C supplemented with 1 mM isoxanthine. Cells were harvested at the logarithmic phase by centrifugation at 5,000 × *g* for 10 min at 4 °C. Total RNA was extracted using a commercial RNA isolation kit, and residual genomic DNA was removed by DNase treatment. RNA quality was assessed, and mRNA was enriched by depleting ribosomal RNA (rRNA). RNA library construction and high-throughput sequencing were performed by Novogene (China). Clean reads were aligned to the reference genome using Bowtie2 (v2.2.3). Gene-level read counts were obtained using HTSeq (v0.6.1)^43^. Differential gene expression analysis between CarBT4gout 2.0 and EcN1917 wild-type strains cultured with isoxanthine was performed using DESeq2. Genes with adjusted *p*-values (Padj) < 0.05 and |log₂ fold change| > 1 were considered significantly differentially expressed and were visualized in a volcano plot^44^.

### Animal husbandry and sample collection

Six-week-old male uricase-deficient (UOX⁻/⁻) mice on a C57BL/6JGpt background (B6/JGpt- Uoxem3Cd3501/Gpt; GemPharmatech, Jiangsu, China) were used. Mice were maintained on standard chow and allopurinol-supplemented water (100 mg/L) until 6 weeks of age, after which they were switched to normal drinking water. Animals had ad libitum access to food and water, and were housed under a 12-h light/dark cycle at 20-22 °C with 40-60% humidity. Bacterial suspensions containing CarBT4gout 2.0 or wild-type EcN1917 (WT) strains were prepared at a density of 1 × 10^10^ cfu per 200 μL PBS. Mice were administered bacteria or PBS vehicle control daily by intragastric gavage over a period of 6 weeks. Blood samples were collected from the caudal vein of live mice into tubes containing concentrated sodium EDTA (final concentration ∼12 mM) and allopurinol (final concentration ∼12 µM) to inhibit xanthine oxidase^45^. Plasma was obtained by centrifugation at 1,500 × *g* for 10 min at 10 °C. All samples were stored at -80 °C until analysis. The animal care and experimental protocols conformed to the recommendations in the 8th Edition of the Guide for the Care and Use of Laboratory Animals of the National Institutes of Health (NIH, revised 2011) and were in accordance with the Animal Management Rules of China (Documentation 55, 2001, Ministry of Health, China). Plasma uric acid, creatinine, and urea levels were measured using commercial assay kits (Nanjing Jiancheng Bioengineering Institute; C012-2-1, C011-2-1, C013-2-1) according to the manufacturer’s instructions.

### Clinical samples

Serum was obtained from 25 gout patients and 43 age- and sex-matched healthy volunteers under protocols approved by the Institutional Review Board of [Tianjin First Central Hospital]. After collection, blood samples were allowed to clot at room temperature for 30 min and centrifuged at 1,500 g for 10 min at 4 °C. Aliquots of serum (50 µL) were deproteinized by addition of 150 µL ice-cold acetonitrile:MeOH (1:1, v/v), vortexed for 1 min and centrifuged at 16,000 g for 10 min. Supernatants were analysed on the Agilent 1260 HPLC–6420 triple-quadrupole platform under the MRM conditions described above. Serum uric acid and isoxanthine concentrations were quantified against external calibration curves.

### Quantification and statistical analysis

All statistical analyses and definitions of sample sizes are provided in the figure legends. Unless otherwise indicated, experiments were performed with at least three biological replicates. Error bars in bar and line graphs represent the standard deviation (s.d.). For each sample, three to five independent experiments were conducted, each with three to five technical replicates.

Statistical significance was assessed using Student’s unpaired t-test. A *p* value < 0.05 was considered statistically significant (* *p* < 0.05, ** *p* < 0.01, *** *p* < 0.001).

### Bioinformatics analysis

To construct the sequence similarity network of the SsnA family (IPR017700) in bacteria, a non-redundant sequence set was assembled by downloading all UniProt entries annotated with the “SsnA family” domain (IPR017700) and mapping them to their corresponding UniRef90 clusters (proteins clustered at ≥ 90 % sequence identity), excluding fragment sequences. An SSN was generated in EFI-EST^46^ using an edge E-value cut-off of ≤ 10⁻¹⁴⁰. Genome- neighbourhood information within a 15-ORF window was retrieved with EFI-GNT^46^ to detect co-occurrence with selected protein families. The network was visualised in Cytoscape^47^.

## Acknowledgments

We thank the instrument analytical center of the School of Pharmaceutical Science and Technology at Tianjin University for providing the LC-MS analysis.

## Author contributions

Z.L., Y.Wei, and Y.Z. conceptualization; Z.L., W.M., Z.G., W.P., J.Z., Y.W., X.W., Z.Z., C.Z., Z.T., Z.N., Y.T., Z.H. and Y.Wei data curation; Z.L., W.M., Z.G., J.Z., Y.Wei, and Y.Z. formal analysis; Y.Wei, and Y.Z. funding acquisition; Z.L., W.M., Z.G.,W.P., J.Z., Y.W., Y.Wei, and Y.Z. methodology; Y.Wei, and Y.Z. project administration; Y.Wei, and Y.Z. resources; Y.W., and Y.Z. supervision; Z.L., W.M., Z.G., W.P., J.Z., Y.Wei, and Y.Z. validation; Z.L., W.M., Z.G, W. P., Y.Wei and Y.Z. visualization; Z.L., W.M. Z.G, W. P., J.Z, Y.Wei, and Y.Z. writing-original draft; Z.L., W.M., Z.G, W. P., Y.Wei, and Y.Z. writing review and editing.

## Data and materials availability

All data needed to evaluate the conclusions in the paper are present in the paper and/or the Extended data.

## Ethics statement

This study was approved by the Animal Research Committee of Shanghai Jiao Tong University (NO. A2024400-003).

---

**Table S1.** Strains used in this study

| Strains | Description | Reference/Source |
| --- | --- | --- |
| *E. coli* MG1655 | Coli Genetic Stock Center strain (CGSC) No. 6300 | CGSC |
| *E. coli* Nissle 1917 | Coli Genetic Stock Center strain | CGSC |
| *E. coli* MG1655 $\Delta xdhA$ | The *xdhA*-deleted mutant of *E. coli* MG1655 | This study |
| *E. coli* MG1655 $\Delta xdhD$ | The *xdhD*-deleted mutant of *E. coli* MG1655 | This study |
| *E. coli* Nissle 1917 $\Delta xdhA$ | The *mdtL*-deleted mutant of *E. coli* Nissle 1917 | This study |
| *E. coli* Nissle 1917 $\Delta xdhD$ | The *mdtG*-deleted mutant of *E. coli* Nissle 1917 | This study |
| *E. coli* Nissle 1917 $\Delta ygfK$ | The *ygfK*-deleted mutant of *E. coli* Nissle 1917 | This study |
| CarBT4gout 2.0 strain | The *pgapA-pnirBp* insertion of *E. coli* Nissle 1917 | This study |
| CarBT4gout 2.0 strain;<br>$\Delta xdhA$ | The *xdhA*-deleted mutant of *E. coli* CarBT4gout 2.0 strain | This study |
| CarBT4gout 2.0 strain;<br>$\Delta xdhD$ | The *xdhD*-deleted mutant of *E. coli* CarBT4gout 2.0 strain | This study |

**Table S2.**
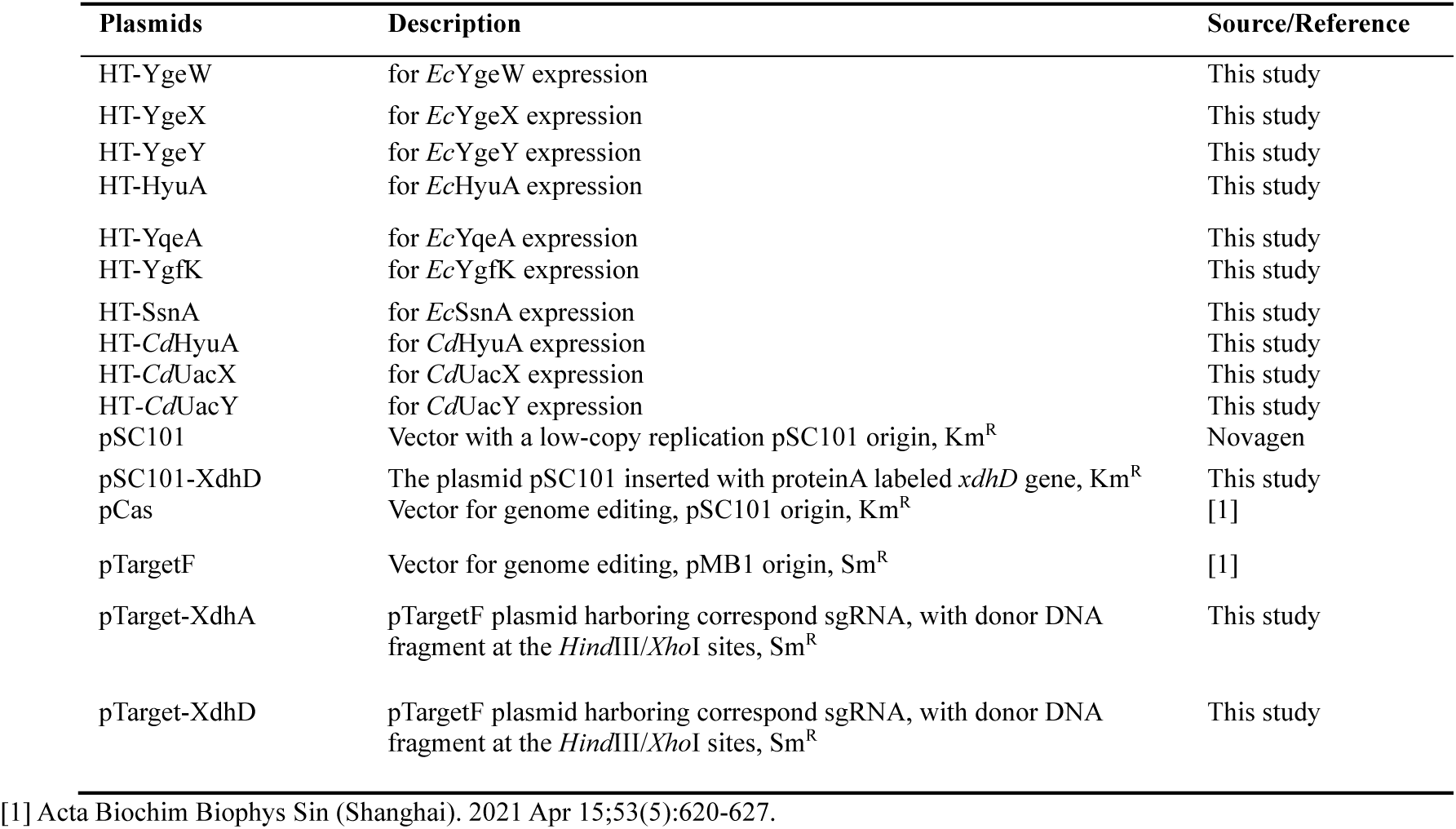
Plasmids used in this study

| Plasmids | Description | Source/Reference |
| --- | --- | --- |
| HT-YgeW | for *EcYgeW* expression | This study |
| HT-YgeX | for *EcYgeX* expression | This study |
| HT-YgeY | for *EcYgeY* expression | This study |
| HT-HyuA | for *EcHyuA* expression | This study |
| HT-YqeA | for *EcYqeA* expression | This study |
| HT-YgfK | for *EcYgfK* expression | This study |
| HT-SsnA | for *EcSsnA* expression | This study |
| HT-*CdHyuA* | for *CdHyuA* expression | This study |
| HT-*CdUacX* | for *CdUacX* expression | This study |
| HT-*CdUacY* | for *CdUacY* expression | This study |
| pSC101 | Vector with a low-copy replication pSC101 origin, Km<sup>R</sup> | Novagen |
| pSC101-XdhD | The plasmid pSC101 inserted with proteinA labeled *xdhD* gene, Km<sup>R</sup> | This study |
| pCas | Vector for genome editing, pSC101 origin, Km<sup>R</sup> | [1] |
| pTargetF | Vector for genome editing, pMB1 origin, Sm<sup>R</sup> | [1] |
| pTarget-XdhA | pTargetF plasmid harboring correspond sgRNA, with donor DNA fragment at the *HindIII/XhoI* sites, Sm<sup>R</sup> | This study |
| pTarget-XdhD | pTargetF plasmid harboring correspond sgRNA, with donor DNA fragment at the *HindIII/XhoI* sites, Sm<sup>R</sup> | This study |

[1] Acta Biochim Biophys Sin (Shanghai). 2021 Apr 15;53(5):620-627.

**Table S3.**
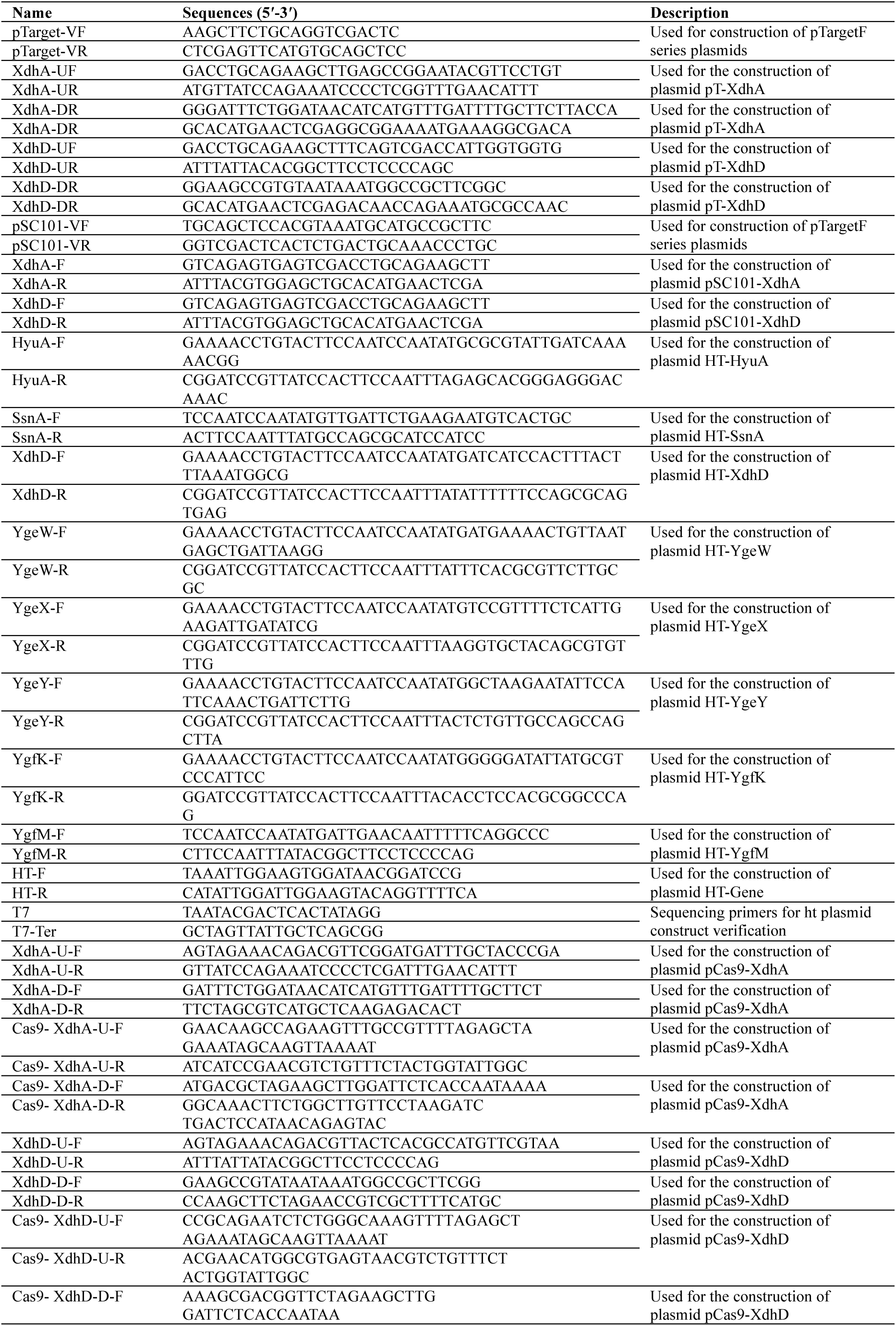
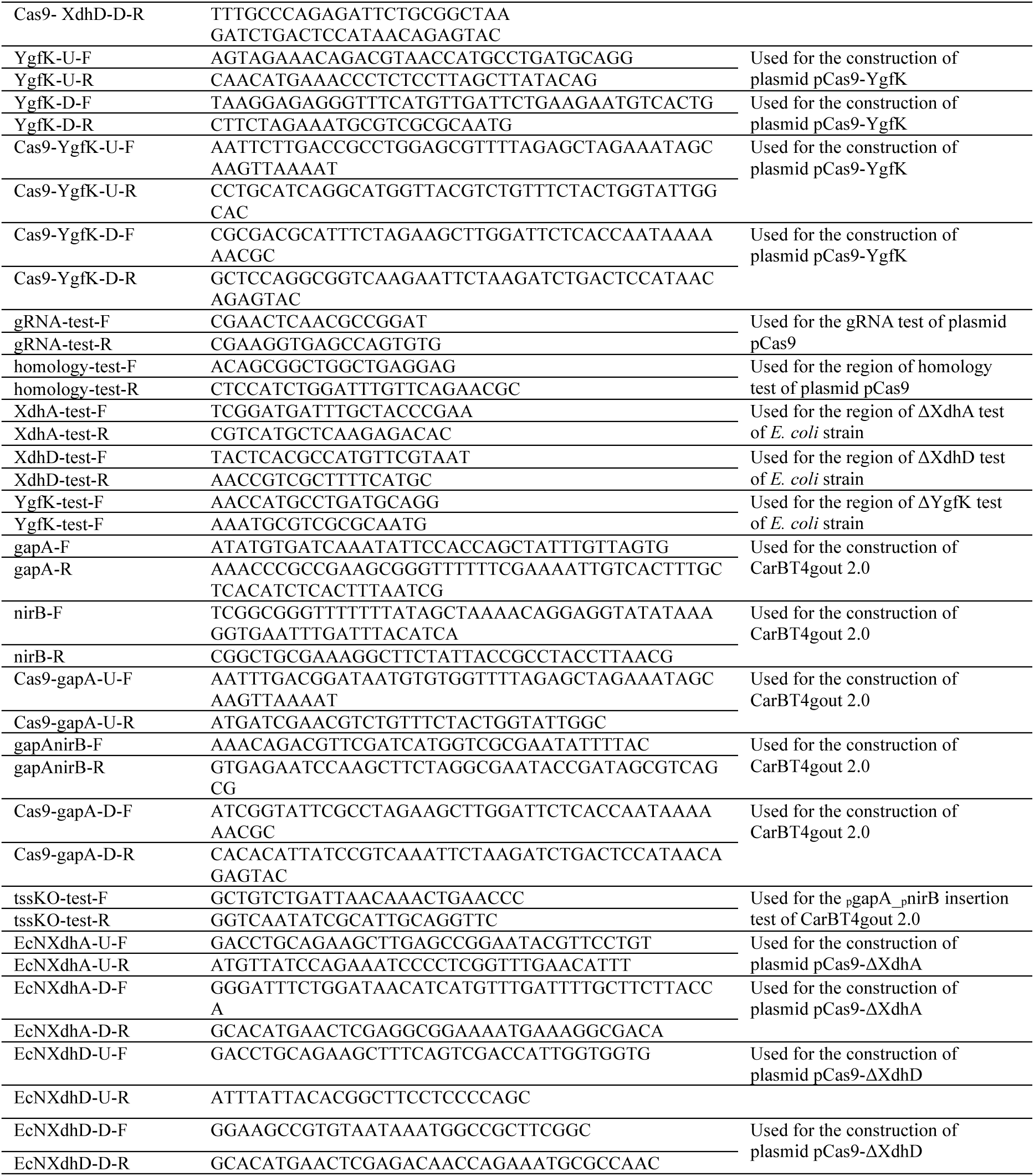
Oligonucleotides used in this study

**Table S4.** Chemicals used in this study.

| Chemicals and Reagents | Source | Identifier |
| --- | --- | --- |
| Acetonitrile, HPLC Grade | MACKLIN | A800362-4L |
| Agar gel strength | Solarbio | A8191-500g |
| Allopurinol | MERYER | M13658-500G |
| Ammonium acetate | MERYER | M22256-100G |
| Ampicillin | BBi | A610028-0025 |
| Calcium chloride dihydrate | Greagent | G10224B |
| Creatinine | Shanghai yuanye Bio-Technology | B25917-100mg |
| Creatinine (N-methyl-D3, 98%) | Shanghai yuanye Bio-Technology | T19668-25mg |
| DTT (Dithiothreitol) | BBi | A620058-0005 |
| Ethanol | MERYER | F30022-4L |
| Glycerol | BBi | A450039-0500 |
| Hypoxanthine | Sangon Biotech | A500336-0050 |
| Kanamycin sulfate | Shanghai yuanye Bio-Technology | S17025-100g |
| L-albizzin | MCE | HY-121167 |
| L-cysteine | BBi | A600132-0100 |
| L-(+)-Arabinose | Adamas | 63128D |
| Lysozyme | Solarbio | L8120 |
| M9 minimal salts | BBi | A507024-0250 |
| Magnesium sulfate heptahydrate | Greagent | G10221C |
| Methanol | MACKLIN | M813903-4L |
| PMSF (phenylmethanesulfonyl fluoride) | Shanghai yuanye Bio-Technology | S30438 |
| Sodium bicarbonate | Aladdin | S112331-500g |
| Sodium chloride | Greagent | G81793J |
| Sodium hydroxide | Aladdin | S111507-500g |
| Spectinomycin dihydrochloride pentahydrate | BBi | A600901-0005 |
| Tris | Shanghai yuanye Bio-Technology |  |
| Tris-HCl | MERYER |  |
| Tryptone | BBi | A650217-0500 |
| Uric acid | MERYER | M23894-25G |
| Uric acid-[1,3-<sup>15</sup>N<sub>2</sub>] | Shanghai yuanye Bio-Technology | B45070-10mg |
| Water | Millipore | Milli-Q® SQ 2 |
| Xanthine | MERYER | M23876-25G |
| Isoxanthine (2,8-dioxopurine) | Shanghai yuanye Bio-Technology | Y90960-1g |
| Yeast extract | OXOID | LP0021B |

**Extended Data Figure 1.**
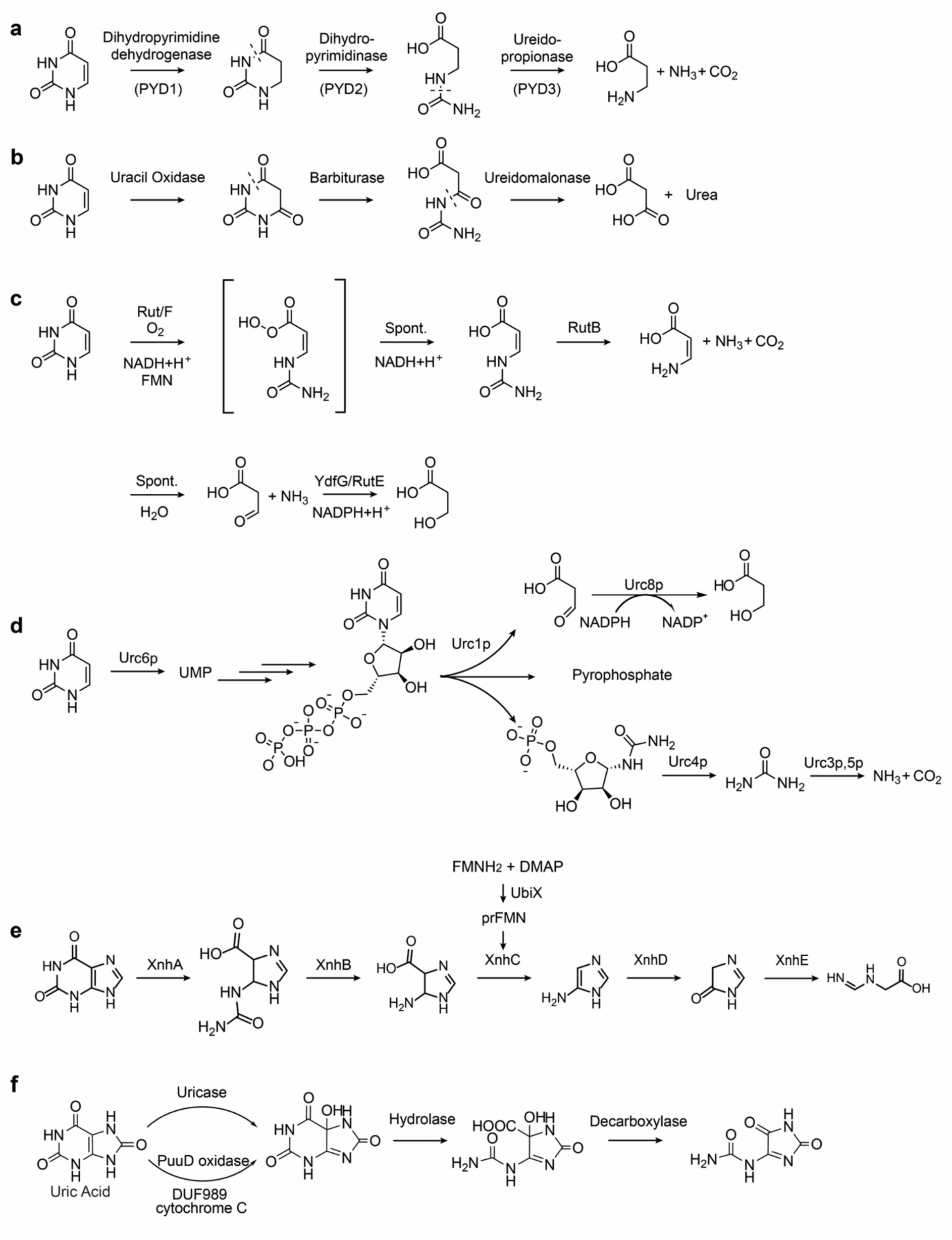
Pyrimidine and purine degradation pathway. a, PYD pathway. **b,** Oxidative pyrimidine degradation pathway. **c,** Rut pathway. **d,** URC pathway in *Lachancea kluyveri.* **e,** Xanthinase pathway. **f,** Oxidative uric acid degradation pathway.

**Extended Data Figure 2.**
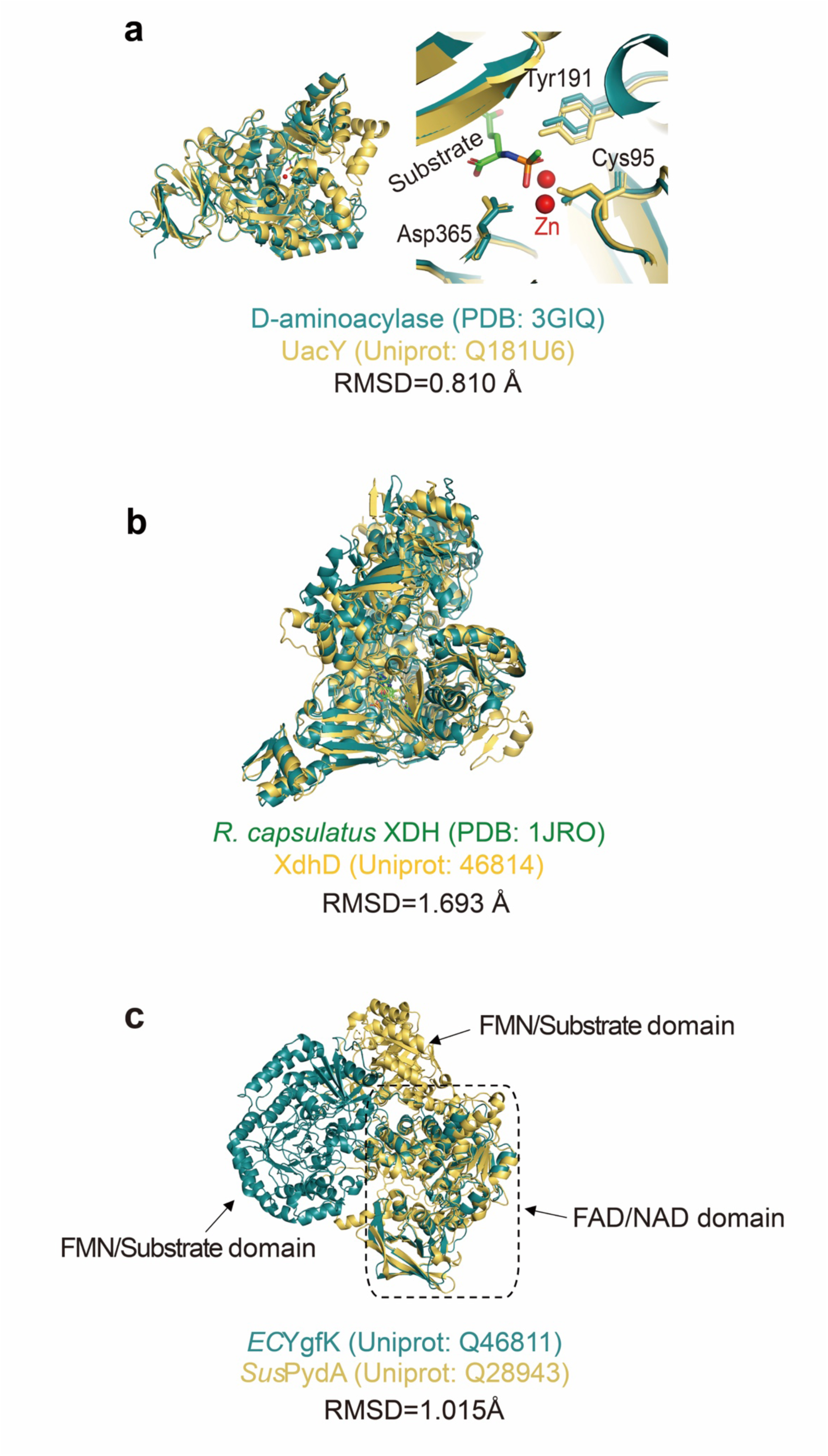
Structural comparisons of UacY, XdhD and YgfK. a, Structures comparisons of D-aminoacylase (PDB:3GIQ) and *Cd*UacY. **b,** Structures comparisons of *Rhodobacter capsulatus* XDH (PDB: 1JRO) and *Ec*XdhD-YgfM. **c,** Structures comparisons of *Ec*YgfK and *Sus*PydA.

**Extended Data Figure 3.**
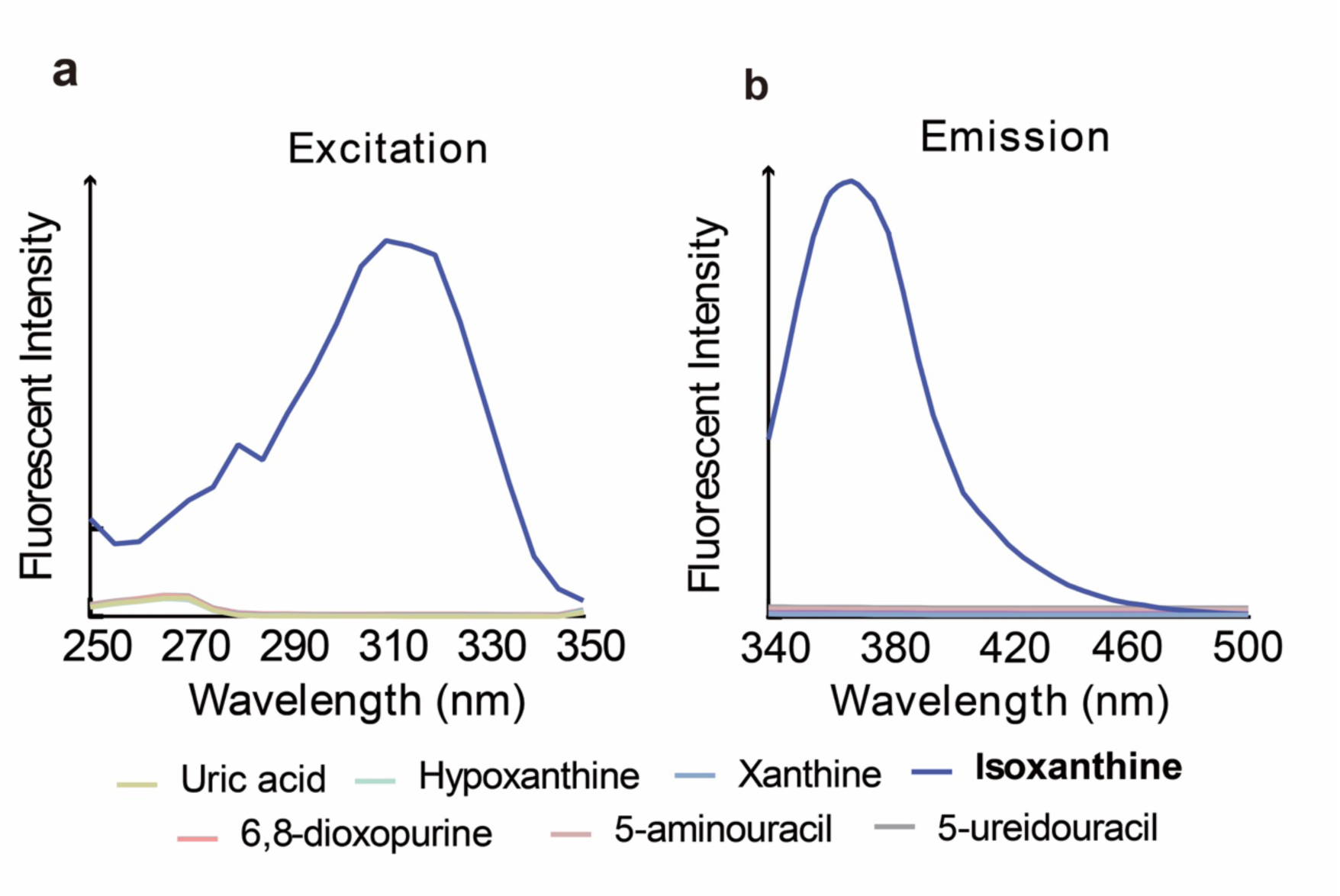
Fluorescence spectrum of purine and pyrimidine derivatives. a, Excitation spectrum. b, Emission spectrum.

**Extended Data Figure 4.**
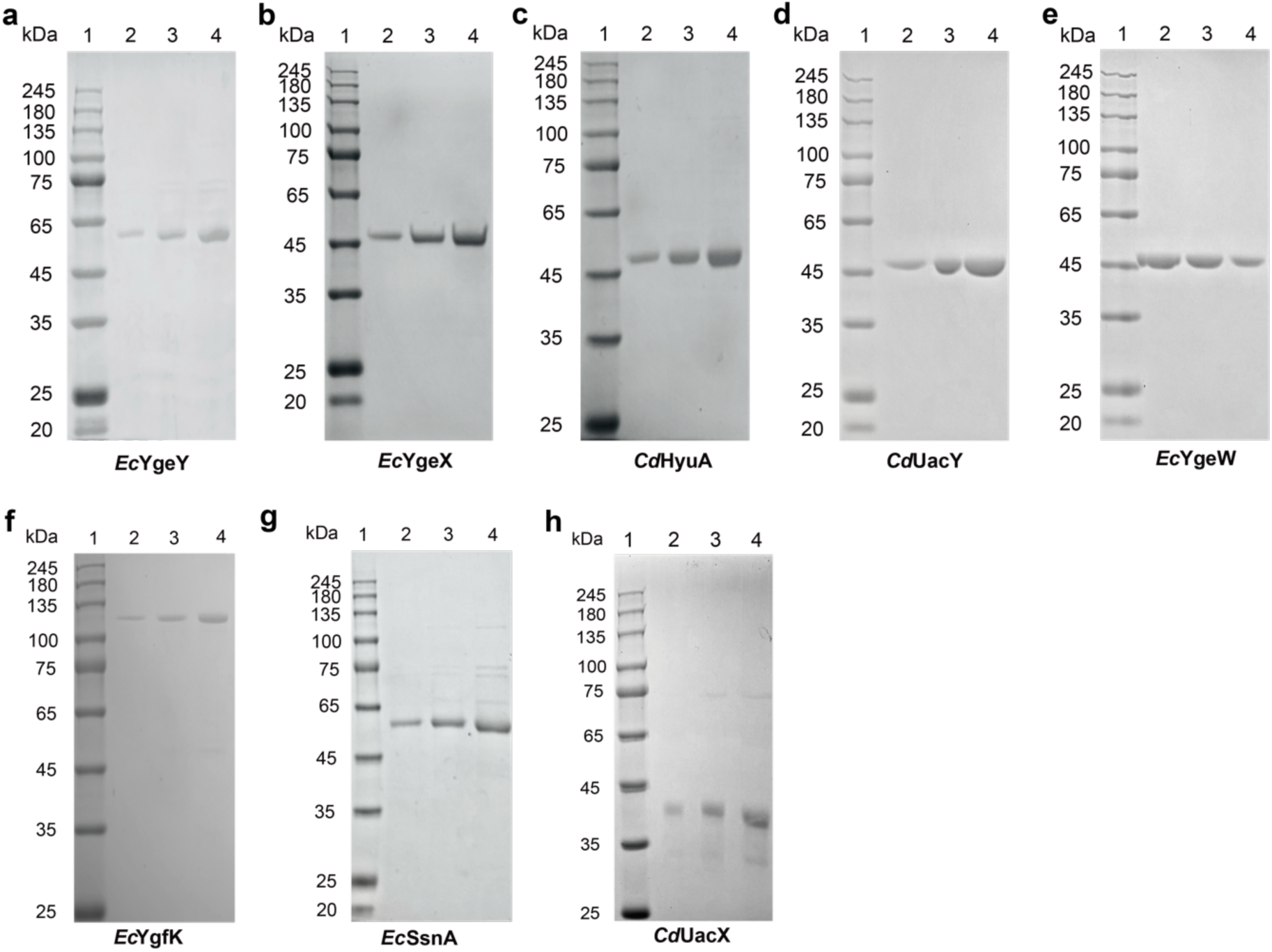
SDS-PAGE analyses of purified proteins used in enzymatic assays and biochemical characterization. a, *Ec*YgeY. b, *Ec*YgeX. c, *Cd*HyuA. d, *Cd*UacY. e, *Ec*YgeW. f, *Ec*YgfK. g. *Ec*SsnA. h, *Cd*UacX. Each 4-20% gradient gel (Bis-Tris) contains protein molecular weight marker and 1, 2, 4 μg of the recombinant protein from lane 1 to lane 4 in order.

**Extended Data Figure 5.**
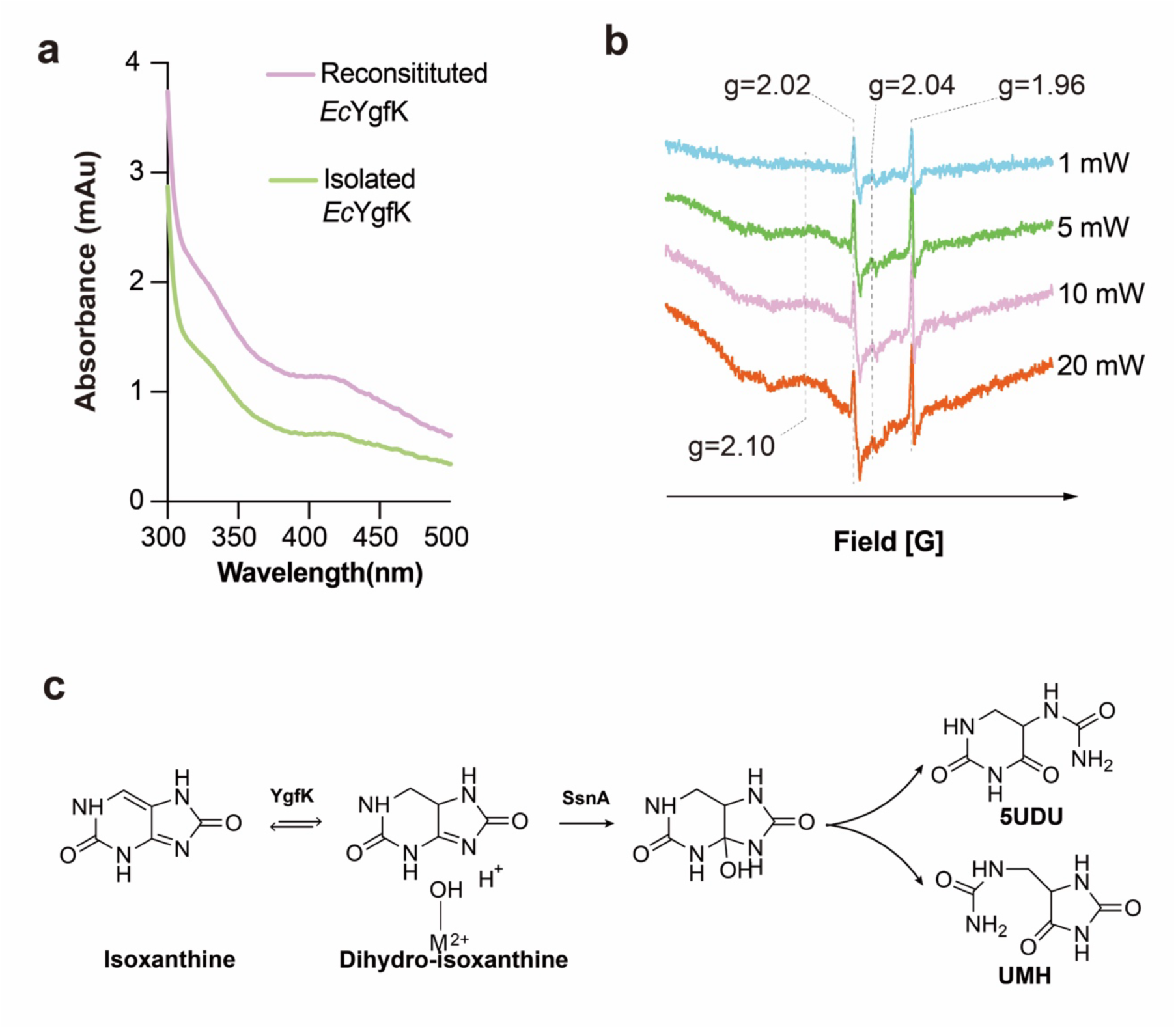
YgfK iron-sulfur cluster and SsnA mechanism. a, UV-Vis absorption spectra of YgfK as isolated and reconstituted. The feature at 410 nm corresponding to [4Fe4S]^2+^ clusters in reconstituted YgfK. **b,** EPR spectrum of reconstituted YgfK. **c.** Proposed mechanism for SsnA.

**Extended Data Figure 6.**
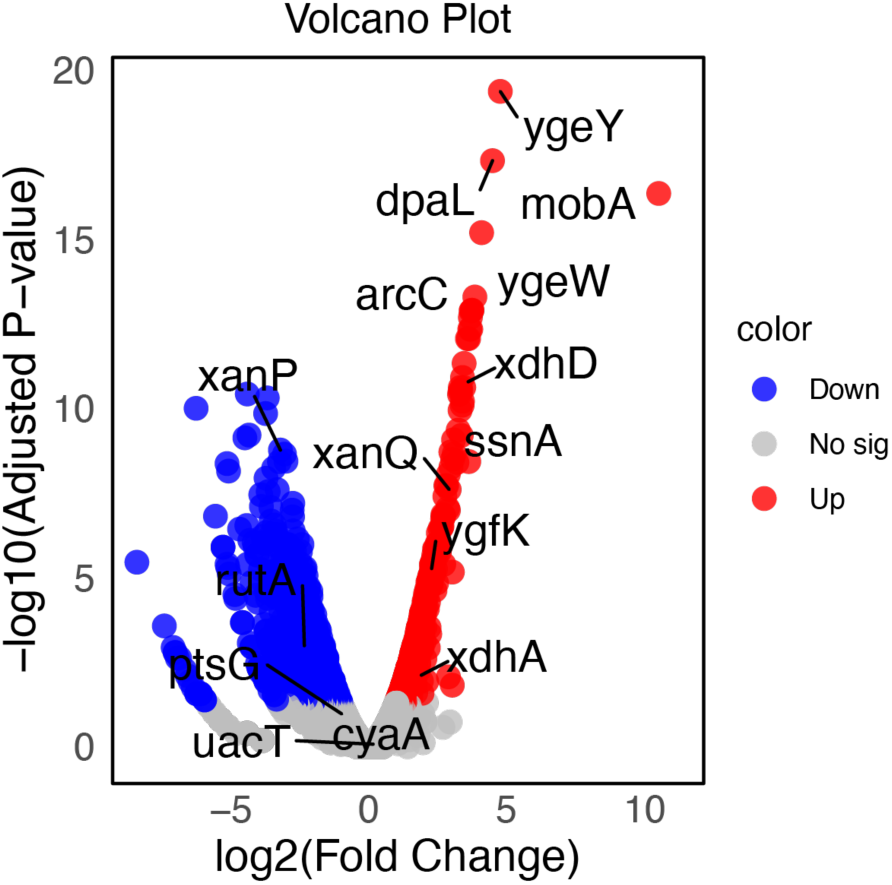
Volcano plot showing differential expression results of uric Acid degradation pathway genes in wide-type EcN1917 and CarBT4gout_2.0 strains. RNA-seq results comparing responder to non-responder biopsies via DESeq2. Differential expression analysis was performed using DESeq2, applying a two-sided Wald test. *p*-values were adjusted for multiple comparisons using the Benjamini–Hochberg method to control the false discovery rate (FDR). Data are shown as Log2 fold changes with associated adjusted *p*- values.

